# A neoantigen-microbead platform for personalized T cell cancer vaccination

**DOI:** 10.1101/2025.11.21.689667

**Authors:** Long Jiang, Birce Akpinar, Yue Li, Maryam Sanaee, Nathaniel Alvin Sanjaya, Kendra Schmidt, Alejandro Fernandez Woodbridge, Ola B Nilsson, Luigi Notari, Csaba Kindler, Ingo Rimke, Olivia Thomas, Shigeaki Kanatani, Per Uhlén, Jerker Widengren, Torbjörn Gräslund, Fredrik Wermeling, Hans Grönlund

## Abstract

Colorectal cancer (CRC) is a leading cause of cancer mortality and is characterized by a high tumor mutational burden, making it responsive to immunotherapy. We developed a bioinformatic and manufacturing pipeline for personalized cancer vaccines and evaluated it in the MC38 colon adenocarcinoma mouse model. Thirty-six high-scoring neoantigens (NAGs) were identified by exome and transcriptome analysis, produced as six purified polypeptides, and coupled to paramagnetic beads. Intralymphatic vaccination of C57BL/6 mice with NAG beads induced robust NAG-specific T cell and antibody responses, resulting in significant inhibition of MC38 tumor growth. Treated tumors displayed increased necrosis and CD8+ T cell infiltration. Compared with soluble peptides, bead-coupled antigens elicited superior protection. Studies in T cell-deficient and antibody-depleted mice confirmed that both CD4+ and CD8+ T cells mediated the antitumor effect. These findings highlight the potential of NAG bead vaccination as an effective immunotherapy for CRC.

## Introduction

The World Health Organization (WHO) estimates that CRC is the third most commonly diagnosed cancer worldwide, with over 1.8 million new cases reported annually^1,2^. Additionally, it ranks as the second leading cause of cancer-related deaths, emphasizing the urgent need for innovation in diagnosis, treatment, and prevention^3^.

Cancer immunotherapy represents a transformative paradigm in the field of oncology, harnessing the inherent capabilities of the immune system to recognize and eradicate malignant cells^4,5^. Unlike traditional cancer treatments that directly target cancer cells, immunotherapy aims to stimulate, modulate, or enhance the body’s immune responses to specifically target and eliminate cancerous cells while minimizing damage to normal tissues^6,7^. Personalized cancer treatment, often synonymous with precision medicine, is revolutionizing cancer care by tailoring therapies to the unique genetic, molecular, and biological characteristics of an individual patient’s cancer^8^. This approach acknowledges the heterogeneity among tumors and the variable responses to standard treatments^9^. The immune system has evolved to distinguish self from non-self, allowing it to identify and eliminate pathogens or abnormal cells. Mutations in cancer cells can produce neoantigens (NAGs), abnormal proteins that the immune system can potentially recognize as tumor-specific targets^10,11^. However, tumors often evade immune detection through various mechanisms, including the expression of checkpoint proteins that inhibit immune responses^12^. Checkpoint inhibitors, such as PD-1 and CTLA-4 inhibitors, help the immune system target these cancer mutations by releasing these inhibitory “brakes”, thereby reactivating T cells to attack the tumor. This has transformed cancer therapy, offering significant clinical benefits, but long-term responses remain inconsistent^13,14^. Cancer vaccines present a promising strategy to enhance the immune system’s ability to target specific mutations. Recent studies, such as those by Moderna and BioNTech, demonstrate the potential of exploiting the cancer mutanome for tumor vaccination^15,16^. These vaccines aim to induce a robust and durable immune response against cancer-specific antigens, potentially improving the efficacy of immunotherapy and providing long-term protection against cancer recurrence^17,18^.

Advances in microbead technology have enabled the development of platforms with tunable sizes and customizable surfaces, facilitating efficient co-delivery of tumor-specific antigens and immune stimulants directly to antigen-presenting cells (APCs). This targeted particulate delivery enhances antigen uptake and T cell activation, promoting robust and sustained anti-tumor immune responses. By concentrating adjuvants and antigens within immune tissues, such platforms may reduce systemic exposure, a concept supported by broader nanoparticle and biomaterial studies, although direct evidence that microbeads specifically decrease off-target effects or systemic toxicity is still limited^19–21^. Our group has previously applied microbead-based antigen presentation across both autoimmune disease and cancer^22,23^. In oncology, this technology has advanced toward clinical translation through the development of personalized tumor-trained lymphocytes (pTTL). We established a bead-based platform enabling the rapid preparation and purification of patient-specific neoantigens, which supports the *ex vivo* expansion of neoantigen-reactive T cells from colorectal cancer patients for subsequent adoptive transfer in an ongoing clinical trial (NEOGAP-CRC-01)^24^.

Here, we explore the potential application of NAG coupled beads as a cancer vaccine within a murine MC38 cancer model. Initially, we selected 36 neoantigens with high scores in MC38 cancer cells. Subsequently, these neoantigens were coupled with microbeads and administered via intralymphatic injection into the inguinal lymph nodes of mice as part of a vaccination strategy. Our findings robustly affirm that the utilization MC38 NAG bead vaccination effectively restrains MC38 tumor growth, and that this inhibition is contingent upon both CD4 and CD8 T cells.

## Results

### Coupling neoantigens identified in MC38 cells to Sera-Mag SpeedBeads

The MC38 cell line is a widely utilized murine model for colorectal carcinoma^25,26^. MC38 has numerous mutations and is sensitive to immunotherapy with checkpoint inhibition^27,28^. To identify mutations and neoantigens, we conducted comprehensive analyses employing both whole exome sequencing (WES) and transcriptome sequencing (RNA seq), and our analysis revealed the presence of over 4000 mutations that were consistently identified in both WES and RNA seq datasets. To rank the mutations, we integrated the allele frequency with the citation number from the Candidate Cancer Gene Database (CCGD)^29^. This scoring system favored mutations demonstrating higher expression levels and greater relevance to cancer. Through this approach, we selected 36 mutations with the highest score (**Sup. Table 1**).

Subsequent to this identification, we formulated six NAG polypeptides, each comprising six NAG peptides (21 mer) separated by a flexible GGS linker of three amino acids, as well as albumin binding domain (ABD) and His-tag (**Fig. 1A** and **Sup. Fig. 1**). These recombinant MC38 NAG polypeptide constructs were synthesized utilizing *Escherichia coli* (**Sup. Fig. 2**) and covalently coupled to paramagnetic Sera-Mag SpeedBeads. These coupled beads, individualized from MC38 cells, are hereby referred to as MC38 beads (**Fig. 1A**). The efficacy of the coupling process was validated through flow cytometry analysis, confirming a high coupling efficiency (**Sup. Fig. 3**).

**Figure 1.**
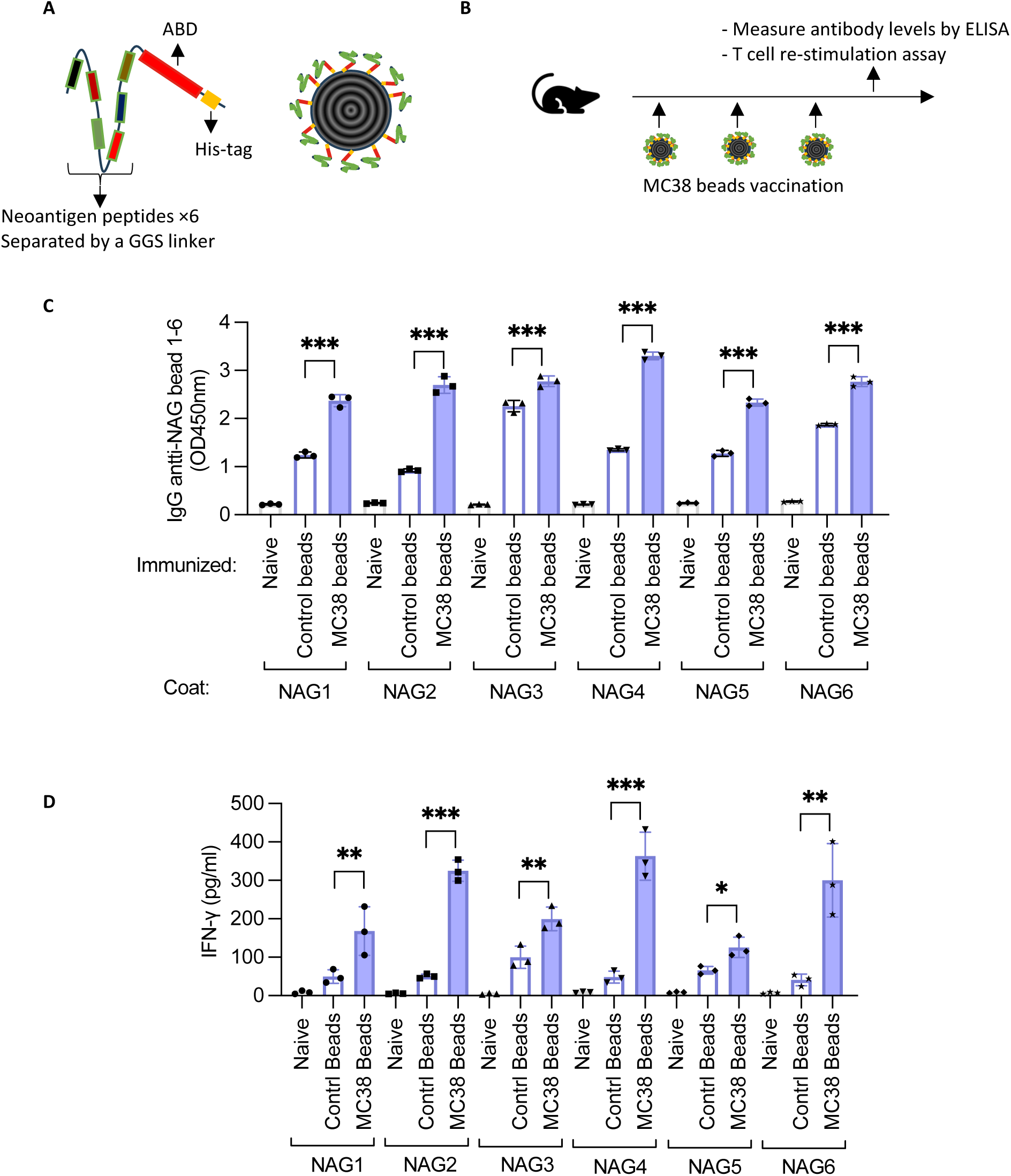
MC38 bead vaccination induces specific immune responses. **A**. Structure of MC38 NAG polypeptides (left) and MC38 beads (right). **B**. Schematic of the mouse vaccination protocol. **C**. Sera were collected from naïve and vaccinated mice (2 weeks after the third vaccination), diluted 1:1600, and analyzed by ELISA on plates coated with NAG1-NAG6 MC38 beads. Optical density (OD) values were measured. **D**. Splenocytes from naïve and vaccinated mice (2 weeks after the third vaccination) were stimulated with NAG1–NAG6 MC38 beads, and IFN-γ levels in the supernatant were measured 2 days later. Data are presented as individual values with mean ± SEM, n = 3 (C-D). Statistical significance: *, p < 0.05; **, p < 0.01; ***, p < 0.001 by one-way ANOVA with Tukey’s post-test.

**Figure 2.**
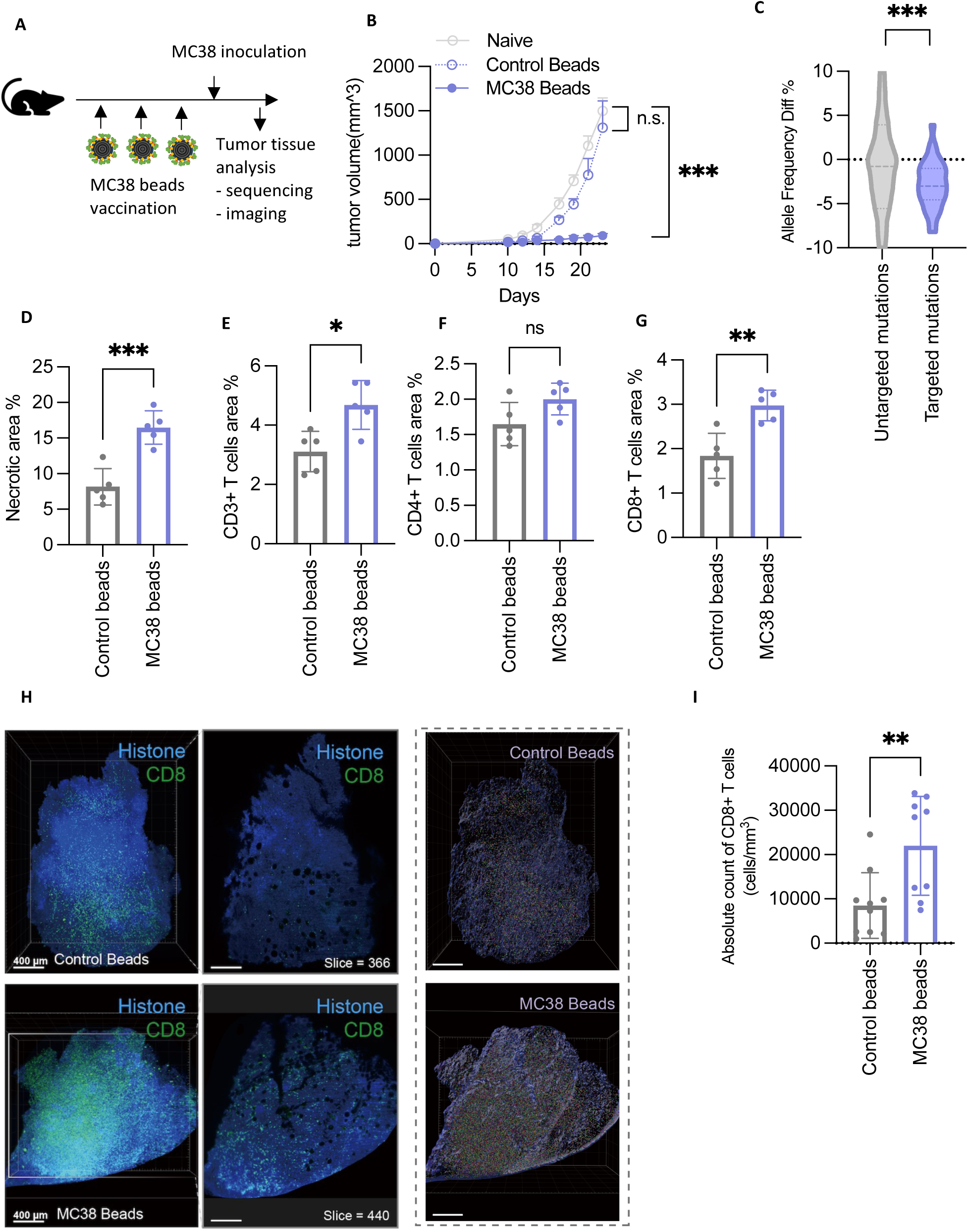
MC38 bead vaccination inhibits MC38 tumor growth. **A**. Schematic of the mouse vaccination and MC38 tumor model setup. **B**. Mice were vaccinated with control beads or MC38 beads, and tumor growth was monitored over time. **C**. Tumors from MC38 bead vaccinated mice were collected at 200–300 mm³ for WES and RNA-seq. Allele frequencies were analyzed and compared with MC38 cells, with differences assessed between bead-targeted and untargeted mutations. **D-G**. Tumors collected at 200–300 mm³ were analyzed histologically. D. H&E staining and quantification of necrotic areas. E-G. Immunohistochemistry for CD3+(E), CD4+ (F), and CD8+ (G) T cells, with areas quantified. **H-I**. Tumors collected at 200–300 mm³ were subjected to 3D imaging. Tissues were stained for Histone and CD8, and CD8+ T cells were quantified. Data are shown as mean ± SEM (B), violin plots (C), or individual values with mean ± SEM (D–G, I). Sample sizes: n = 5 (B, D–G), n = 9 (I). Statistical significance: ns, not significant; *, p < 0.05; **, p < 0.01; ***, p < 0.001 by two-way ANOVA with Tukey post-test for the last time point (B), Welch’s t test (C), and unpaired t test (D–G, I).

**Figure 3.**
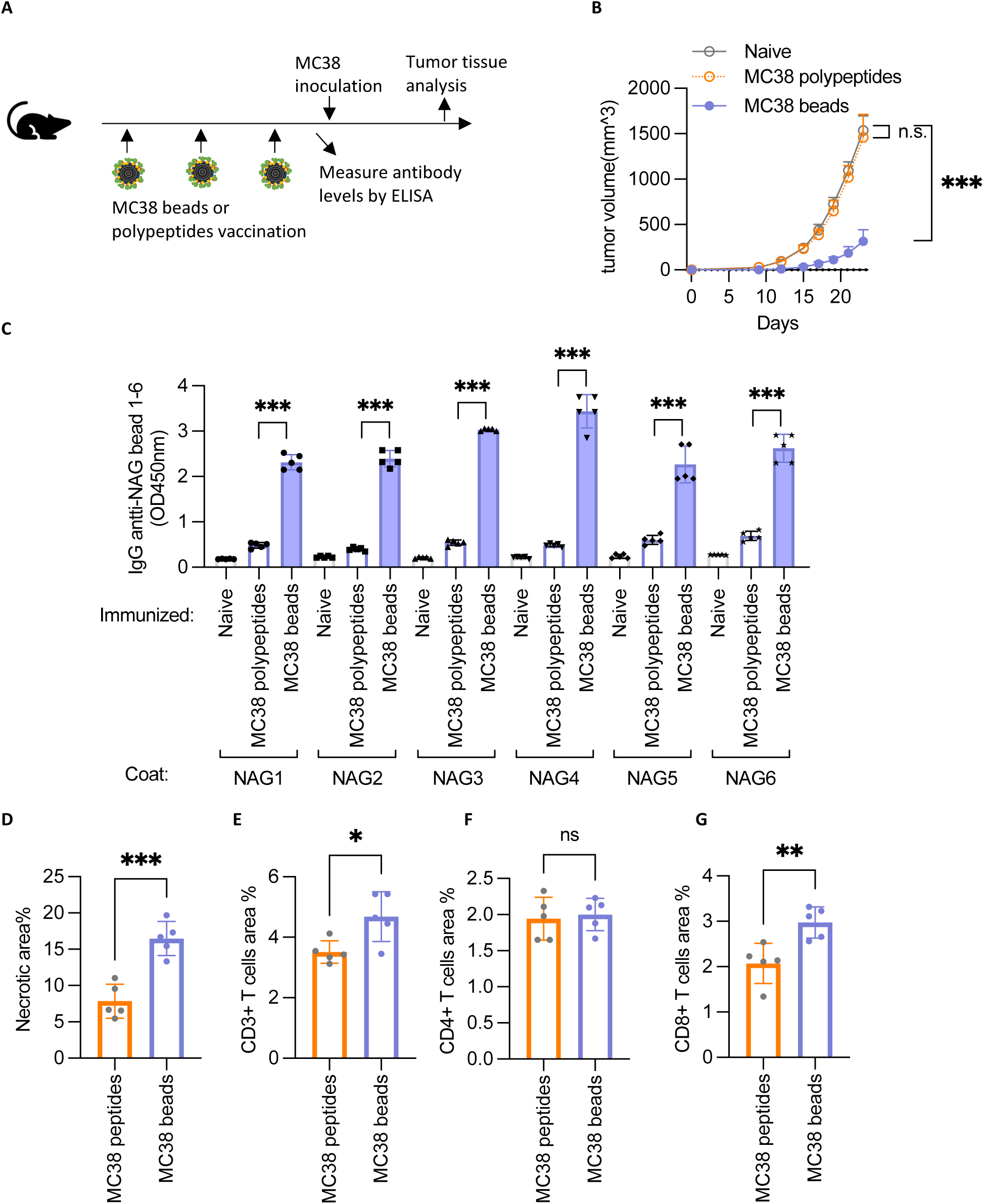
MC38 beads elicit stronger immune responses than soluble polypeptides. **A**. Schematic of the mouse vaccination and MC38 tumor model setup. **B**. Mice were vaccinated with MC38 polypeptides or MC38 beads, and tumor growth was monitored over time. **C**. Sera were collected from naïve mice and from vaccinated mice 2 weeks after the third vaccination, diluted 1:1600, and analyzed by ELISA on plates coated with NAG1–NAG6 MC38 beads. Optical density (OD) values were measured. **D-G**. Tumors were harvested at 200–300 mm³ for histological analysis. D. H&E staining and quantification of necrotic areas. E-G. Immunohistochemistry for CD3+ (E), CD4+ (F), and CD8+ (G) T cells, with areas quantified. Data are presented as mean ± SEM (B) or individual values with mean ± SEM (C-G), n = 5.. Statistical significance: ns, not significant; *, p < 0.05; **, p < 0.01; ***, p < 0.001 by two-way ANOVA with Tukey post-test for the last time point (B), one-way ANOVA with Tukey post-test (C), and unpaired t test (D–G).

### MC38 bead vaccination induces specific immune responses

After we successfully coupled MC38 beads, we investigated the ability of MC38 bead vaccination to induce a targeted immune response in C57BL/6 mice. To maximize the efficacy of the immunization process, we employed intralymphatic injection into the inguinal lymph nodes^30^. The vaccination schedule consisted of three administrations of MC38 beads spaced two weeks apart, as outlined in **Fig. 1B**.

We found that the vaccinated mice developed robust NAG-specific antibodies after MC38 beads vaccination (**Fig. 1C** and **Sup. Fig. 4**). NAG-specific T cells were detected after the third vaccination, as demonstrated by IFN-γ secretion (**Fig. 1D**) and by TNF-α FluoroSpot assay (**Sup. Fig. 5**) following *ex vivo* stimulation of splenocytes with MC38 beads. Both T cell re-stimulation assays revealed a consistent pattern: specific T cells responded to all NAG1- NAG6 beads, with varying magnitudes of response - for example, a stronger response to NAG4 and a weaker response to NAG5.

**Figure 4.**
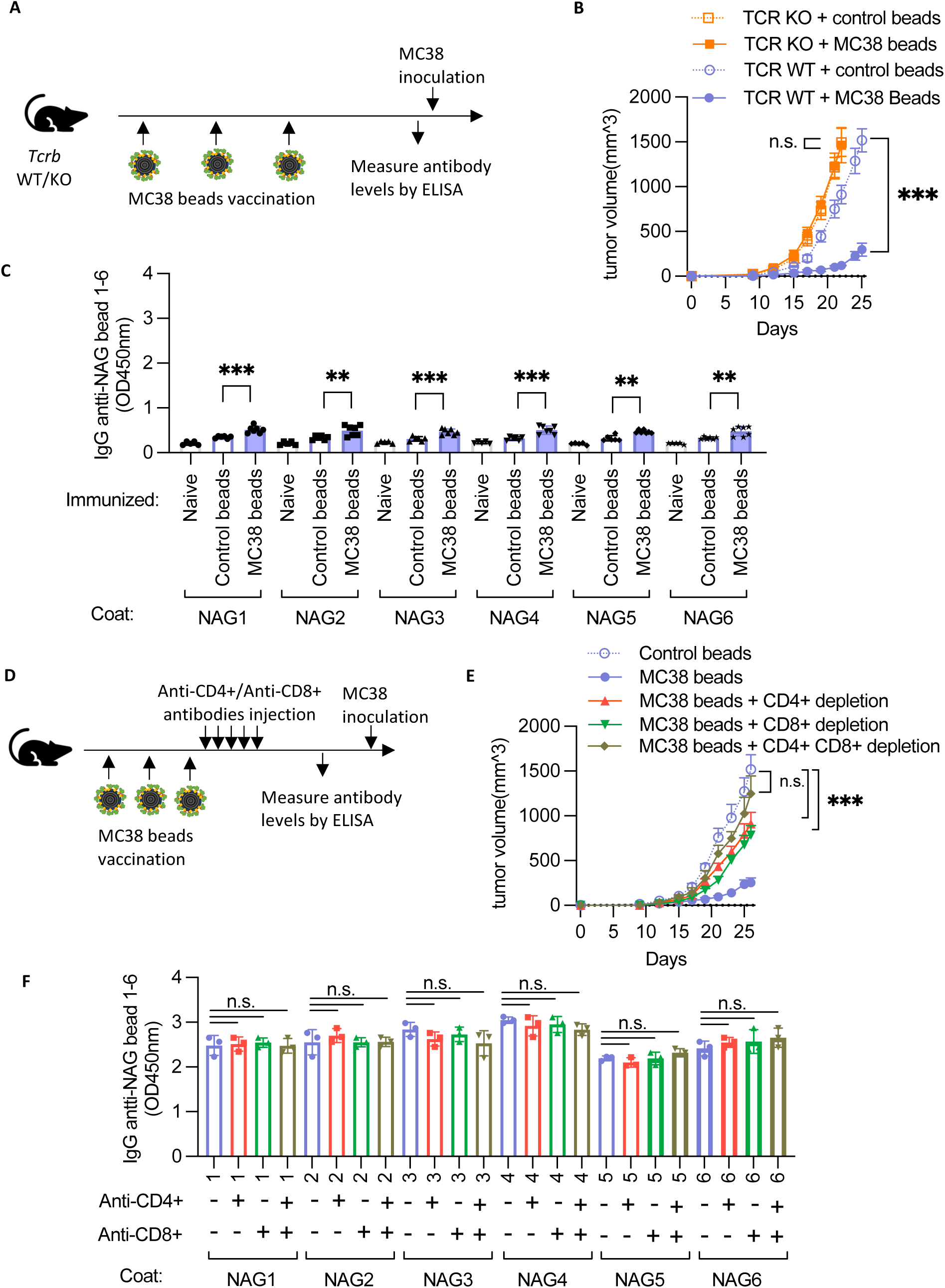
MC38 bead vaccination inhibits tumor growth in a T cell–dependent manner. **A**. Schematic of the mouse vaccination and MC38 tumor model setup. **B**. *Tcrb* wild-type (WT) or knockout (KO) mice were vaccinated with control beads or MC38 beads, and tumor growth was monitored over time. **C**. Sera were collected from naïve TCR KO mice and from vaccinated TCR KO mice 2 weeks after the third vaccination, diluted 1:1600, and analyzed by ELISA on plates coated with NAG1-NAG6 MC38 beads. Optical density (OD) values were measured. **D**. Schematic of the mouse vaccination, T cell depletion, and MC38 tumor model setup. **E**. Mice were vaccinated with control or MC38 beads, with CD4+, CD8+, or combined CD4+/CD8+ T cells depleted before tumor inoculation, and tumor growth was monitored over time. **F**. Sera were collected from MC38 bead-vaccinated mice and from T cell depleted mice prior to MC38 tumor inoculation, diluted 1:1600, and analyzed by ELISA on plates coated with NAG1-NAG6 MC38 beads. OD values were measured. Data are presented as mean ± SEM (B, E) or individual values with mean ± SEM (C, F), n = 5–7 (B, C) or n = 3 (E, F). Statistical significance: ns, not significant; *, p < 0.05; **, p < 0.01; ***, p < 0.001 by two-way ANOVA with Tukey post-test for the last time point (B, E), and one-way ANOVA with Tukey post-test (C, F).

**Figure 5.**
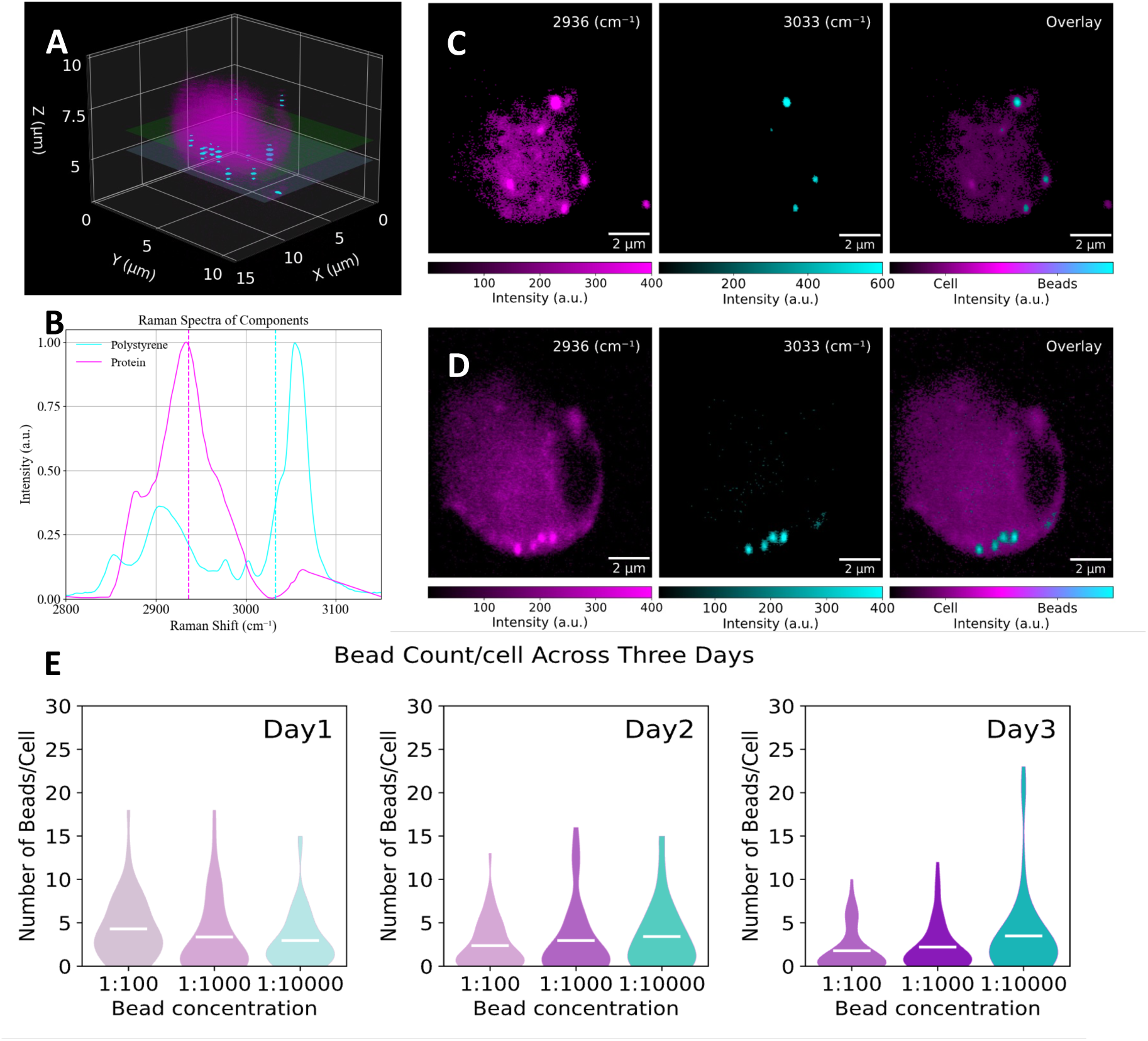
3D SRS imaging of antigen-presenting cells (APCs) and internalized NAG-coated beads. **A**. 3D representative APC generated from multiple XY sections using SRS microscopy. **B**. Normalized Raman spectra of cellular protein (magenta) and polystyrene beads (cyan), with dashed vertical lines marking the wavenumbers selected for imaging. **C-D**. Sectional 2D images of the same APC shown in (A), acquired at 2936 cm⁻¹ (protein, magenta) and 3033 cm⁻¹ (beads, cyan), with overlays illustrating five beads in (C) and four beads in (D). **E**. The overall results of statistics on beads’ uptake per cell through 50 cells per three different beads concentrations and three; white lines indicate mean values.

In summary, MC38 bead vaccination effectively induced a targeted immune response, engaging both humoral (NAG-reactive antibodies) and cellular (NAG-specific T cells) immunity.

### MC38 bead vaccination inhibits MC38 tumor growth

To assess whether the MC38-specific immune response could effectively inhibit MC38 tumor cells, we subcutaneously introduced MC38 cells to the vaccinated mice, as illustrated in **Fig. 2A**. In line with the triggered immune responses, tumor growth was noticeably restrained in mice subjected to MC38 bead vaccination (**Fig. 2B**).

Next, we conducted WES and RNA seq of the tumors in vaccinated mice to explore whether the vaccination applied any selection pressure on the targeted neoantigens. The analysis revealed a significant reduction in the allele frequency of the 36 targeted mutations compared to untargeted mutations (**Fig. 2C**). This indicates that the vaccination applies a selection pressure on the tumor cells expressing the targeted mutations, potentially contributing to its anti-tumor efficacy.

Histological analysis of tumor tissues revealed increased necrosis in vaccinated tumors (**Fig. 2D**), which was associated with enhanced T cell infiltration, particularly CD8⁺ T cells (**Fig. 2E-G**). Three-dimensional imaging further confirmed elevated CD8⁺ T cell infiltration in vaccinated tumors (**Fig. 2H-I** and **Sup. Fig. 6A**) and revealed distinct “hot spots” - areas within the tumor microenvironment characterized by high immune cell activity^31^. In vaccinated mice, these hot spots displayed reduced blood vessel density compared with controls, alongside increased CD8⁺ T cell infiltration (**Sup. Fig. 6B-F**). This pattern suggests that vaccination not only enhances immune cell infiltration into regions of active tumor by immune interaction but may also remodel the vasculature in a way that creates a less favorable environment for tumor progression.

In conclusion, MC38 bead vaccination suppressed MC38 tumor growth by enhancing CD8⁺ T-cell infiltration and applying selective pressure against tumor cells carrying the targeted neoantigens, thereby reshaping the tumor microenvironment in favor of anti-tumor immunity.

### MC38 beads elicit stronger immune responses than soluble polypeptides

While previous studies have utilized NAG polypeptides for vaccination^32,33^, it remains unclear whether MC38 bead vaccination can elicit stronger immune responses than soluble NAG polypeptides. To ensure a fair comparison, we assumed 100% coupling efficiency for the beads so that the total amount of NAG polypeptides on the beads did not exceed that of the corresponding soluble polypeptide group (**Fig. 3A**). Consistent with enhanced uptake by antigen-presenting cells, MC38 bead vaccination effectively inhibited MC38 tumor growth, whereas soluble NAG polypeptides showed no measurable effect (**Fig. 3B**). Moreover, ELISA analysis revealed that soluble NAG polypeptides induced weaker antibody responses compared with bead vaccination (**Fig. 3C** and **Sup. Fig. 7**). Histological examination of tumors from mice receiving soluble NAG polypeptides showed reduced necrosis and diminished T-cell infiltration relative to bead-vaccinated tumors (**Fig. 3D-G**).

In summary, these results highlight the importance of using MC38 beads as a neoantigen carrier, as bead vaccination induces more potent immune responses and effectively suppresses MC38 tumor growth.

### MC38 bead vaccination inhibits tumor growth in a T cell-dependent manner

So far, we have demonstrated the significant inhibition of MC38 tumor progression by bead vaccination, coupled with increased T cell infiltration within the tumor tissue. Building upon these observations, we next investigated whether the inhibitory effect relies on the presence of T cells using α/β T cell-deficient *Tcrb* knockout (KO) mice in our study, as outlined in **Fig. 4A**.

As anticipated, MC38 bead vaccination failed to protect *Tcrb* KO mice, indicating that the vaccine’s anti-tumor effect requires α/β T cells (**Fig. 4B**). Notably, MC38 tumors grew faster in *Tcrb* KO mice than in wild-type controls, demonstrating that endogenous α/β T cells suppress MC38 growth even in the absence of vaccination. These observations are consistent with the well-characterized immunogenicity of the MC38 model - an immune-inflamed (“hot”) colorectal tumor model that is sensitive to T cell–mediated control and to immune checkpoint interventions^34,35^. This observation was further supported by ELISA data, revealing that vaccination failed to elicit robust antibody titers in *Tcrb* KO mice (**Fig. 4C**), in stark contrast to the response observed in *Tcrb* WT mice (**Fig 1C**). Interestingly, the MC38 beads still induced some antibody production in *Tcrb* KO mice. This indicates that while the bulk of the antibody response is T cell-dependent, a certain extent of the response could also be T cell-independent, perhaps due to the repetitive nature of the bead structure^36^.

To identify the T cell subsets responsible, we depleted CD4+ and/or CD8+ T cells in bead-vaccinated mice prior to MC38 tumor transfer (**Fig. 4D**). Depletion of either CD4+ or CD8+ T cells alone partially reduced tumor inhibition, whereas combined depletion of both subsets completely abolished the vaccine-induced tumor suppression (**Fig. 4E**). Notably, when we measured antibody levels by ELISA just before tumor transfer, we still observed very high titers of vaccine-specific antibodies - comparable to those in Fig. 1C - and no significant difference in antibody levels following CD4+ or CD8+ T cell depletion (**Fig. 4F**). Combined with the data in Fig. 4E, this indicates that although the vaccine elicits strong antibody responses, these antibodies are not sufficient to suppress tumor growth and serve primarily as a biomarker for successful vaccination.

Together, these findings demonstrate that the anti-tumor effect of MC38 bead vaccination is T cell-dependent, with both CD4+ and CD8+ T cells playing essential roles, while the vaccine-induced antibodies reflect vaccination success rather than contributing to tumor suppression.

### Fate of NAG-coated beads in antigen-presenting cells tracked by 3D stimulated Ramen scattering microscopy

Next, we investigated the fate of NAG-coated beads within APCs was tracked by using label-free 3D stimulated Raman scattering (SRS) microscopy ^37–41^, with subcellular resolution. Purified mouse dendritic cells were co-cultured with varying dilutions of MC38 beads for different time periods, followed by fixation for SRS microscopy analysis. The SRS-3D image of a typical APC is shown in XYZ coordinates (**Fig. 5A**), with two representative sections depicting the intracellular bead distribution (**Fig. 5C** and **Fig. 5D**), showing five and four beads within the cell, respectively. The cell nucleus appears as a dark hole in the SRS images after thresholding due to the distinct peak of Raman shifts for DNA compared to proteins^37^. The beads, visualized as bright spheres in the overlay images (**Fig. 5C** and **Fig. 5D**), are distinguished by the SRS signal of only the polystyrene core at 3033 cm⁻¹ and combination of polystyrene and the NAG-protein coating at 2936 cm⁻¹ wavenumbers.

Tracking the NAG-coated beads by 3D-SRS imaging in 50 cells per sample, multiple beads were found in the APCs, primarily localized at the membrane or cytoplasm of the cells. Only one cell showed a single bead in the nucleus. The bead sizes, measured via the full width at half maximum (FWHM) of corresponding line profiles (**Sup. Fig. 9**), were broadly distributed between 400 and 1000 nm. The distribution of bead numbers per APC was analyzed for three bead concentrations over up to three days of co-culturing (**Fig. 5E**), with the summary of these bead counts per cell statistics (Mean±SD) presented in **Sup. Table 2**. On Day 1, the distribution of number of beads per cell correlated with the bead concentration, with higher bead concentrations resulting in a higher bead uptake by the cells. However, for Day 2 and Day 3, this trend reversed, showing higher bead counts per cell at lower concentrations. This suggests that APCs with higher bead concentrations (1:100 dilution) did not survive as long, while those with fewer beads (1:10000 dilution) exhibited a gradual increase in bead uptake over time. On Day 3, cells incubated with lower bead concentrations showed a statistically significant increase in their bead uptake with longer incubation times.

In conclusion, 3D-SRS microscopy effectively tracked the fate of NAG-coated beads in APCs. The mean number of beads per cell ranged from 2 to 4 across all bead concentrations and incubation times. The majority of beads were localized to the membrane and cytoplasm, with minimal bead presence in the nucleus.

## Discussion

In this study, we investigated the efficacy of personalized neoantigen-coupled beads as a vaccine platform for inducing protective immune responses against cancer neoantigens in the MC38 murine colorectal adenocarcinoma model. Our findings highlight several critical insights into the mechanisms underlying the observed anti-tumor effects and the potential of bead-based vaccination strategies.

For NAG selection, we applied a straightforward prioritization strategy - integrating allele frequency from WES and RNA-seq data with gene relevance scores from the CCGD citation data - to identify 36 top-ranking mutations. While this approach did not incorporate the full spectrum of modern neoantigen prediction parameters, such as peptide-MHC binding affinity or antigen processing likelihood, it nevertheless yielded a panel that elicited strong and protective immune responses in the MC38 model. The robust outcome from this relatively simple selection method suggests that, at least in certain tumor models, high-expression and cancer-relevance criteria may capture a substantial fraction of immunogenic targets. This finding is encouraging for the development of rapid, scalable vaccine pipelines, particularly in settings where time or resources limit the use of more complex prediction algorithms. While effective, our findings suggest that factors beyond expression levels and cancer relevance - such as antigen presentation efficiency and T cell recognition - may further refine neoantigen selection^42^. Future improvements in predictive algorithms could enhance the precision of this step, optimizing vaccine efficacy^43^.

We chose a prophylactic vaccination approach, introducing MC38 cells after establishing an antigen-specific immune response. This design allowed us to evaluate the maximal potential of the vaccine-induced immunity to prevent tumor establishment, minimizing the confounding effects of the rapid tumor growth kinetics characteristic of the MC38 model. While this setup demonstrates the capacity of the neoantigen-coupled bead platform to elicit potent and protective T cell responses, it does not fully address its therapeutic efficacy in established tumors. Given that checkpoint blockade, particularly anti-PD-1 therapy^44^, has shown substantial efficacy in the MC38 model due to its high mutational burden and immunogenicity, future studies should explore whether combining neoantigen-coupled bead vaccination with PD-1 inhibition can generate synergistic antitumor activity in a therapeutic setting. Such combination strategies could better model clinical scenarios, where treatment often begins after tumor detection.

A noteworthy discovery of our study was the direct comparison between MC38 NAG-coupled beads and soluble NAG polypeptides. By ensuring antigen dose equivalence (assuming 100 % coupling efficiency), we observed that bead-based vaccination markedly outperformed soluble polypeptides in restraining tumor growth. This finding aligns with growing evidence that particulate antigen delivery platforms significantly outperform soluble peptide vaccines in activating the immune system^45,46^.

The critical role of T cells in MC38 bead-mediated tumor inhibition was confirmed using *Tcrb* KO mice, which lack functional α/β T cells. Vaccination failed to provide tumor protection in these mice, and tumor growth was accelerated compared to wild-type controls, highlighting the necessity of T cell-mediated immunity. Interestingly, residual antibody production observed in *Tcrb* KO mice suggests that some aspects of the humoral response may occur independently of T cells, potentially due to the repetitive bead structure that can directly engage B cells.

Further experiments with selective depletion of CD4+ and CD8+ T cells demonstrated that both subsets contribute to the anti-tumor effects of MC38 bead vaccination. While the depletion of either subset alone reduced the protective activity of the vaccine, the complete abrogation of tumor inhibition following double depletion highlights the collaborative role of CD4+ and CD8+ T cells. These findings underscore the importance of bead-bound antigens in facilitating antigen cross-presentation, which drives effective T cell activation and coordination.

A distinctive strength of our study is the application of label-free 3D SRS microscopy to directly visualize the fate of NAG-coated beads inside APCs with subcellular resolution. Unlike conventional fluorescence approaches that require extrinsic labeling and may alter particle behavior, SRS microscopy allowed us to track these vaccine carriers in their native state, in real time, and in three dimensions. This capability is particularly valuable for particulate vaccine platforms, where understanding intracellular trafficking is central to predicting and optimizing immunogenicity.

Beads were efficiently internalized and localized mainly to membrane and cytoplasmic regions consistent with endo-lysosomal processing, supporting their role in MHC antigen presentation. Beads persisted for days at lower doses, suggesting sustained antigen availability, while heavy loading reduced APC survival. These findings link bead trafficking to the strong immune responses observed and demonstrate SRS microscopy as a powerful, non-invasive tool for guiding particulate vaccine design.

In conclusion, our study establishes MC38 beads as a highly effective cancer vaccine platform that activates robust T cell responses, modulates the tumor microenvironment, and inhibits tumor growth. These findings advance our understanding of the interplay between neoantigen presentation, immune activation, and tumor dynamics. Future research should focus on elucidating the molecular mechanisms driving bead vaccine efficacy and exploring their application across diverse tumor models and clinical contexts. This work represents a significant step toward the development of targeted cancer immunotherapies, offering a promising avenue for enhancing precision oncology.

## Methods

### Cells

The MC38 cell line was purchased from Kerafast (ENH204-FP) and used at a low passage number. The cells were cultured in RPMI-1640 with 10% serum and 1% penicillin-streptomycin-glutamine. Cells were confirmed Mycoplasma negative before culture.

### Animals

Eight- to 12- week-old, sex- and age-matched mice were used for experiments. All animal experiments have been approved by the local animal ethical board at Karolinska Institutet (ethical permit 10681-2020). Mice were housed under specific pathogen-free conditions in a 12-h/12-h light-dark cycle with standard diet ad libitum. WT C57BL/6J, and TCRb KO (B6.129P2-*Tcrb^tm1Mom^*/J) mice were purchased from Charles River Laboratories.

### Cancer Model

1.5×10^6 million cells were injected s.c. on the right flank of the animals, and the tumor size was followed over time by caliper measurement. For bead vaccination, NAG beads (9.5µl) and CpG (0.5µl, 0.05nmol) were used for intralymphatic injection (first two doses) or subcutaneous injection (last dose) close to the injected lymph node. Intralymphatic injection was performed under anesthesia by making a small incision in the inguinal region to expose the inguinal lymph node, followed by direct delivery of the vaccine suspension into the lymph node using a fine gauge micro syringe under visual guidance^47^. Anti-CD4 (clone GK1.5) and anti-CD8 (clone 2.43) (both from BioXcell) were injected i.p. (100µg) 4 times (day -4, -3, -2, and -1) before tumor cell injection (day 0).

### Next-Generation Sequencing and Bioinformatic Analysis

Whole-exome sequencing (WES) and RNA sequencing (RNA-seq) were performed to comprehensively profile tumor mutational and transcriptional landscapes. DNA and RNA were extracted from tumor samples, and WES libraries were sequenced to a mean target coverage of ≥100×, while RNA-seq libraries (poly(A)-selected) were sequenced to a depth exceeding 50 million paired-end reads per sample.

Raw RNA-seq reads were aligned to the *Mus musculus* GRCm39 reference genome using STAR (v2.7) in two-pass mode^48^. Post-alignment, unstranded gene-level read counts were generated by STAR and collated for all samples using a custom R (v4.2) script. Variant calling was performed using VarDict-Java with parameters optimized for accurate detection in RNA-seq data^49^. The expressed allele frequency (AF) reported by VarDict-Java was extracted for downstream analysis. Variants with read depth ≥20 and AF ≥0.4 were retained as high-confidence calls.

Somatic mutations were defined as variants present in both WES and RNA-seq datasets. To rank variants, the allele frequency was integrated with citation counts for each gene from the Candidate Cancer Gene Database (CCGD)^29^, producing a composite score that favored mutations with high expression and established cancer relevance.

### Design, Expression, and Purification of NAGs

The NAG1-6 genes, including flanking BsaI sites, were designed and subcloned into the pET28 vector ^20^ containing an albumin binding domain (ABD)^50^ via a one-step digestion-ligation reaction using BsaI and T4 DNA ligase^51^. Each NAG protein consisted of ABD, six different 21-mer peptides seperated by GGS linkers, and a 6X His-tag at the N-terminal. An ABD protein with a 6X His-tag was used as a negative control (**Sup. Fig. 1**). The expression vectors were transformed into *E. coli* BL21-AI cells (Thermo Fisher Scientific), cultured in super broth medium with kanamycin for 6 hours at 37°C, followed by transfer to auto-induction medium for overnight growth at 25°C. Cells were harvested by centrifugation, and the pellet was frozen. For protein purification, the frozen pellet was thawed with lysis buffer (6 M Guanidinium-HCl, 50 mM Bicine, 10 mM NaCl, 20 mM β-mercaptoethanol, 0.05% Poloxamer, pH 8.0) and sonicated. The lysate was centrifuged, and the supernatant was applied to His Mag Sepharose Ni Beads (Cytiva) purification^20,52^. The proteins were eluted at pH 2.0, the pH was adjusted to 5, and the samples were stored at 4°C. Protein purity and concentration were assessed by SDS-PAGE and NanoDrop. (**Sup. Fig. 2**)

### Coupling of NAGs to paramagnetic beads

NAGs were coupled to Sera-Mag SpeedBeads (Cytiva) via a mine coupling as previously described^20,21^. Briefly, the carboxyl groups on the surface of the beads were activated with 0.1M *N*-hydroxysuccinimide (NHS) and 0.4M 1-ethyl-3-(3-dimethylaminopropyl) carbodiimide (EDC) before the addition of NAGs. After 30 minutes of incubation at room temperature with constant shaking, empty carboxyl groups were deactivated with bicine. Beads were reconstituted in 0.1% Poloxamer containing PBS after the endotoxin washes in a 2M NaOH solution.

The coupling efficacy of each NAG to beads was determined using a monoclonal anti-6X His IgG, CF 488A antibody (Sigma-Aldrich). Beads diluted 1:10 were incubated with the 1:400 diluted antibody for 20 min at RT. After 3 washes with 0,1% Poloxamer containing PBS, volume was adjusted to 150 μl and analysed by flow cytometry (FACSVerse, BD biosciences, USA). Data analysis was performed in FlowJo 10 software (FlowJo LLC, USA) by using each unstained NAG-coupled beads and activated/deactivated beads with no antigen as negative controls. (Sup Figure3)

### ELISA

For assays with splenocytes from vaccinated mice, single-cell suspensions were generated using 40 μm cell strainers (Fisher Scientific), and red blood cells lysed using RBC lysis buffer (ThermoFisher Scientific) following the suggested protocol. Cells were plated in a U-bottom 96-well plate at a concentration of 2×10^5^ cells/well in cRPMI and stimulated as indicated in the figures. IFN- γ secretion was assessed by ELISA according to manufacturer instructions (BioLegend).

To compare the antibody titer towards specific NAG, an in-house ELISA was performed using sera from control and NAGs vaccinated mice. The plates were coated overnight at 4°C with 5 μg/ml recombinant NAG1, 2, 3, 4, 5, 6 or bovine serum albumin (BSA) proteins in 50mM carbonate/bicarbonate buffer, pH 9.6. After 1 hour blocking with PBS containing 1% BSA and 0.05% Tween 20, sera diluted in PBS containing 0.2% BSA and 0.05% Tween 20 were added and incubated for 2 hours which was followed by 1 hour incubation with peroxidase-conjugated AffiniPure goat anti-mouse IgG (Jackson ImmunoResearch). Finally, plates were developed with 3,3′,5,5′-tetramethylbenzidine (TMB) substrate and the reactions were stopped after 10 min with 0.5M H2SO4. Optical density (OD) at 450nm was determined using a SpectraMax Plus 384 using the SoftMax Pro 7.1 software. All tests were performed in duplicate. Pooled sera from vaccinated mice sera, no secondary, and no antigen coating were used as controls. Each mouse’s OD in the BSA wells was subtracted from the NAG wells before statistical analysis. Between each step, plates were washed (4x) with PBS containing 0.05% Tween 20 using ELISA plate washer. All incubation steps were performed at room temperature (RT) unless otherwise stated.

### Fluorospot

Splenocytes from NAG- or ABD-coupled beads vaccinated and non-vaccinated mice were processed into single-cell suspensions by gentle mechanical dissociation through 40 µm cell strainers (Fisher Scientific) in cold medium. Red blood cells were removed by treatment with RBC lysis buffer (Thermo Fisher Scientific) according to the manufacturer’s instructions. Following lysis and washing, splenocytes were counted using an automated cell counter (LUNA-II, Logos Biosystems, South Korea). 150,000 viable cells were seeded in AIM V Medium, AlbuMAX supplement (Gibco) containing 0,05% β-mercaptoethanol to a precoated and blocked TNFalfa FluoroSpot plate (Mabtech) containing NAG or ABD-coupled beads. The plates were incubated and developed according to the manufacturer’s instructions. The FluoroSpot plates were read using an IRIS plate reader (Mabtech) and the SpotReader v.1.1.9 software (Mabtech). Vaccinated mice group’s response to ABD-coupled beads was subtracted from the spot count in the NAG-coupled wells and data are presented as delta Spot Forming Units (ΔSFUs).

### H&E staining and IHC staining

For H&E, formalin-fixed, paraffin-embedded tissue sections (4 µm thick) were performed using standard protocol. For IHC, the sections were deparaffinized and rehydrated, followed by antigen retrieval in citrate buffer (10 mM, pH 6.0) at 95°C for 20 minutes. After blocking endogenous peroxidase with 3% hydrogen peroxide and non-specific binding with 5% BSA, primary antibodies for mouse CD3 (1:200, Abcam, ab16669), CD4(1:4000, Abcam, ab183685), and CD8 (1:4000, Abcam, ab209775) were applied and incubated overnight at 4°C. Sections were then treated with biotinylated secondary antibodies and ABC solution (Abcam), stained with DAB (Abcam), counterstained with hematoxylin, dehydrated, cleared, and mounted. Negative controls omitted the primary antibody.

### 3D imaging

Tumor samples were fixed in 4% PFA overnight at 4°C, followed by methanol dehydration and storage at -20°C. Samples were clarified using 66% DCM in methanol, bleached in 5% hydrogen peroxide in methanol, rehydrated through methanol gradients, and permeabilized in 20% DMSO and 2.3% glycine in PBS with 0.2% Triton X-100 (PTx.2) at 37°C for 24 hours.

For iDISCO+ whole-mount immunostaining, samples were blocked in 6% normal donkey serum and 10% DMSO in PTx.2 for 24 hours at 37°C, incubated with primary antibodies (anti-CD8, Abcam, 1:200; anti-CD31, Abcam 1:200) in PTwH for 3 days at 37°C, followed by washing and secondary antibody incubation for 3 days with Alexa Fluor™-labeled antibodies. Nuclear staining was performed using Histone nanobodies conjugated to Atto 488.

Cleared tumors were embedded in 3% gelatin, dehydrated with methanol gradients, and clarified with DCM and dibenzyl ether (DBE). Imaging was performed using the LaVision Biotec UltraMicroscope II with a z-step size of 2 µm and voxel size 0.755×0.755×2 µm at 488 nm, 561 nm, and 647 nm wavelengths. Images were processed using Imaris software (v.10.2.0, Oxford Instruments) for alignment, intensity normalization, noise reduction, and segmentation, as described previously^41^.

### Stimulated Raman scattering (SRS) microscopy

A schematic of the SRS microscope is shown in Sup Fig 8. The system includes a picoEmerald FT laser (APE GmbH) generating two 2 ps-pulsed beams: a Stokes beam (1032 nm, modulated at 20 MHz) and a tuneable pump beam (660–1010 nm, modulated at 40 MHz). The beams were focused onto cell samples using an oil immersion objective (Olympus, UPLXAPO, 100×, NA = 1.45) and collected in transmission mode with an oil condenser (Olympus, NA = 1.4). The transmitted light was spectrally filtered (ZET980 short pass) and the SRS signal detected in the stimulated Raman loss mode using a Si-photodiode and a lock-in amplifier (Dual frequency SRS detector set, MW Elektooptik). A total of 50 single cells were imaged randomly over the coverslip, with the entire volume of each cell captured in three dimensions using a Z-range of 10–12 μm and Z-steps of 250 nm, alongside an XY-pixel size of 50 nm. Imaging was performed at two selected Raman wavenumbers (**Fig. 5**).

Sera-Mag Carboxylate-Modified Magnetic SpeedBeads (Cytiva), contains polystyrene layers with Raman signals above 3000 cm⁻¹. Based on Raman spectra (**Fig. 5B**)^37,38^, beads and cells were imaged at 3033 cm⁻¹ and 2936 cm⁻¹, corresponding to pump wavelengths of 786 nm and 792 nm, respectively. These were selected to minimize spectral overlap between the SRS signals from the beads and from the proteins and lipids (their CH stretching band) in the cells.

To isolate murine dendritic cells (DCs), murine spleen, inguinal, and axillary lymph nodes were made into single cells suspension by filtering through 0.45 µm membrane. Red blood cells were lysed from the spleen sample by performing ACK lysis (Gibco). Pan mouse dendritic cell isolation kit (Miltenyi) was used to isolate DCs from the single cell suspension, following manufacturer instructions and confirmed with flow cytometry analysis using Live/Dead aqua dead cell staining kit (Invitrogen), CD45.2 (BioLegend, Clone: 104), CD11c (BD Bioscience, Clone: HL3) and MHCII (BioLegend, Clone: M5/114.15.2).

Isolated DCs (A total of 100,000 per treatment group) were co-cultured on coverslips (#1.5, 22 × 22 mm) together with NAG-coated beads at three different beads dilutions (1:100, 1:1000 and 1:10,000) and incubated for up to three days. Co-cultured cells were fixed at the end of the incubation period using 2% paraformaldehyde (Thermo Fisher Scientific) and mounted in PBS using an image spacer (GBL654006, Merck) and then sealed with a coverslip (#1.5, 24 × 60 mm) for SRS imaging.

## Authors’ Contributions

**Long Jiang**: Conceptualization, formal analysis, investigation, visualization, methodology, writing-original draft, writing-review and editing. **Birce Akpinar**: Formal analysis, investigation, visualization, methodology, writing-review and editing. **Yue Li**: Formal analysis, visualization, methodology, writing-review and editing. **Maryam Sanaee**: Formal analysis, visualization, methodology, writing-review and editing. **Nathaniel Alvin Sanjaya**: Visualization, methodology, writing-review and editing. **Kendra Schmidt**: Methodology, writing-review and editing. **Alejandro Fernandez Woodbridge**: Methodology, writing-review and editing. **Ola B Nilsson**: Methodology, writing-review and editing. **Luigi Notari**: Methodology, writing-review and editing. **Csaba Kindler**: Methodology, writing–review and editing. **Ingo Rimke:** Methodology, writing–review and editing. **Olivia Thomas**: Methodology, writing-review and editing. **Shigeaki Kanatani:** Methodology, writing-review and editing. **Per Uhlén**: Funding acquisition, methodology, writing-review and editing. **Jerker Widengren**: Funding acquisition, methodology, writing-review and editing. **Torbjörn Gräslund**: Methodology, writing-review and editing. **Fredrik Wermeling**: Conceptualization, resources, supervision, investigation, writing–review and editing. **Hans Grönlund**: Conceptualization, resources, supervision, funding acquisition, investigation, writing–review and editing, project administration.

## Data Access Statement

The datasets generated during and/or analyzed during the current study are available from the corresponding author on reasonable request.

## Acknowledgement

This study was supported by grants from Cancer- och allergifonden, European Innovation Council and Eurostars. FW received funding from grants provided by the Karolinska Institutet, the Swedish Research Council, and the Swedish Cancer Society. The 3D imaging experiments were supported by the Swedish Research Council (2021-03108, P.U.); the Swedish Brain Foundation (FO2024-0057-TK-138, P.U.); and the Swedish Cancer Society (22 2454 Pj, P.U.). The SRS imaging part of this study (MS, IR and JW) was supported by the European Union’s Horizon 2020 research and innovation program under grant agreement 101017180.

