## Supplementary Figures for "A neoantigen-microbead platform for personalized T cell cancer vaccination"

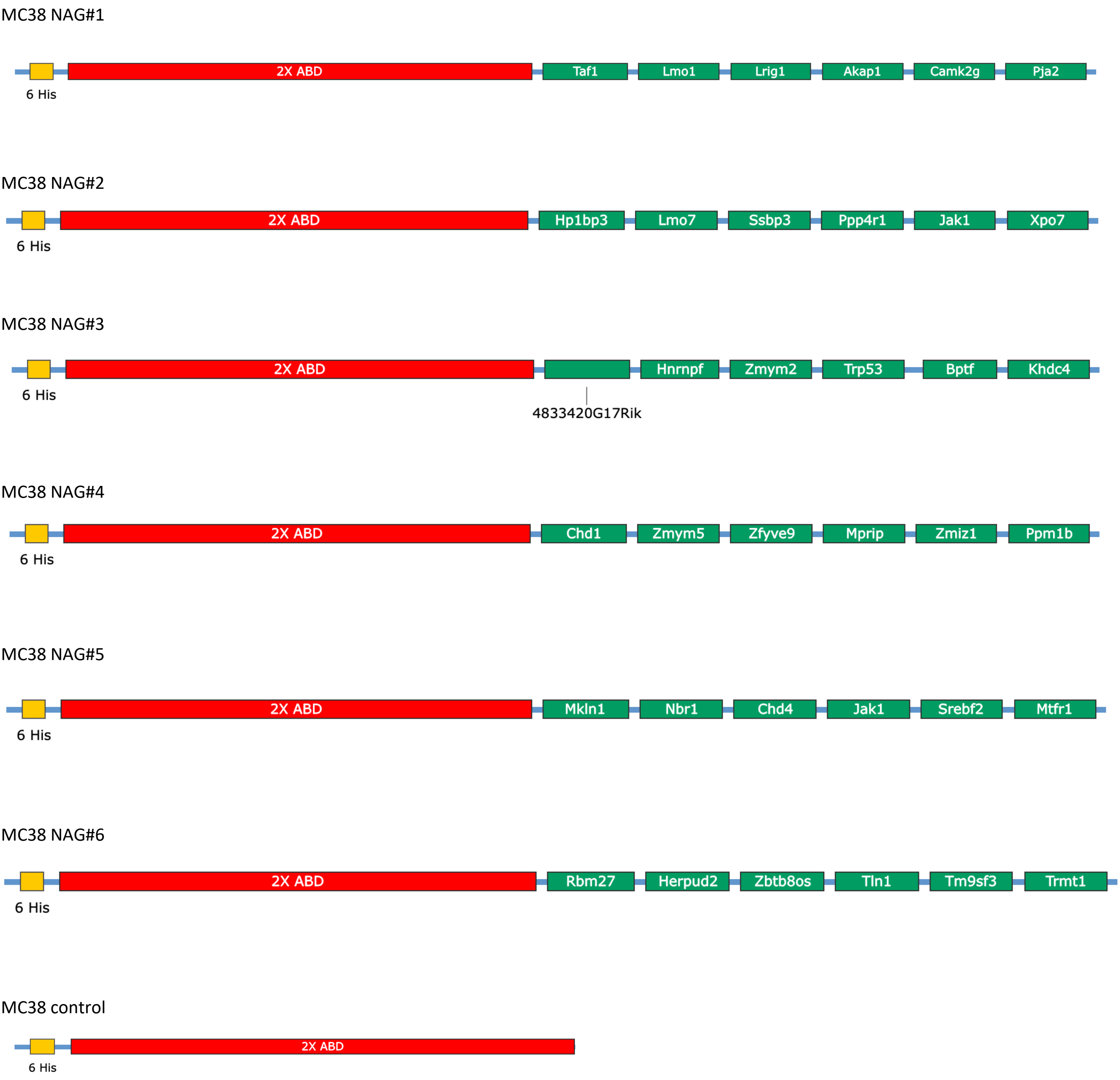


**Supplementary Figure 1.** Schematic of MC38 NAG polypeptides (NAG#1 - NAG#6) and MC38 control polypeptide. Neoantigen polypeptides (NAG#1–NAG#6) consist of six 21-amino-acid peptides (with the point mutation in the middle) linked via GGS linkers and include a 6×His tag and a 2×ABD (albumin-binding domain). The MC38 control construct contains only the 6×His tag and 2×ABD without any neoantigen sequences.

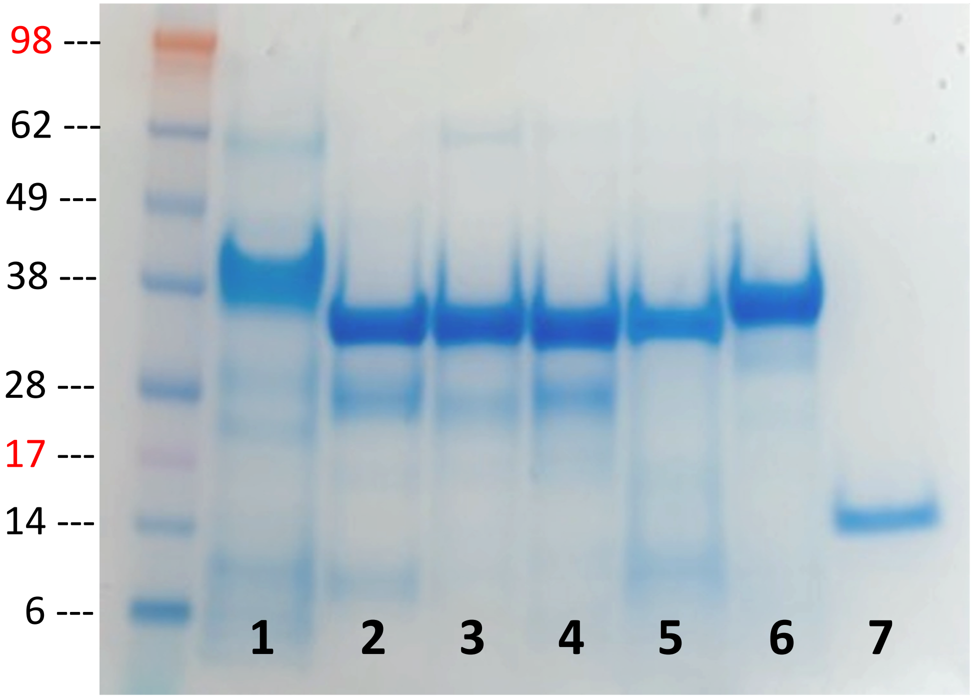


1. NAG1 polypeptide (30.0kDa)
2. NAG2 polypeptide (30.5kDa)
3. NAG3 polypeptide (30.2kDa)
4. NAG4 polypeptide (29.8kDa)
5. NAG5 polypeptide (29.6kDa)
6. NAG6 polypeptide (29.9kDa)
7. Control polypeptide (15.0kDa)

**Supplementary Figure 2,** SDS-PAGE of purified MC38 NAG1-6 and MC38 control polypeptides.

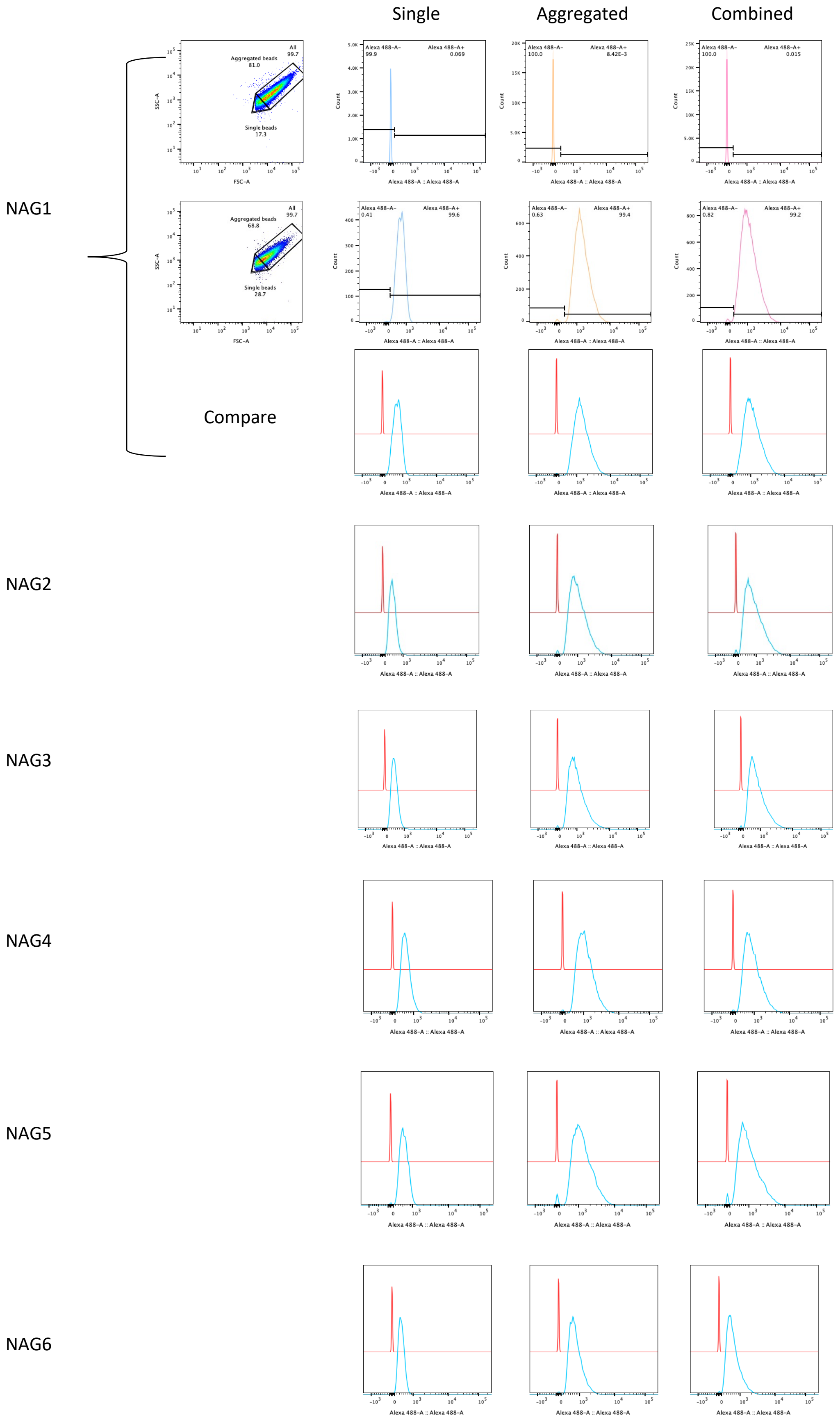


**Supplementary Figure 3**. Flow cytometry analysis of MC38 beads to assess coupling efficiency. MC38 beads coupled with His-tagged NAG1–6 polypeptides were either stained or left unstained with anti–His-tag Alexa Fluor 488 antibody and analyzed by flow cytometry. Data are shown separately for single beads, aggregated beads, and the combined bead population.

A

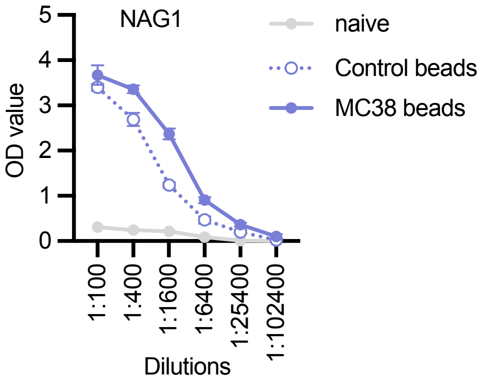


B

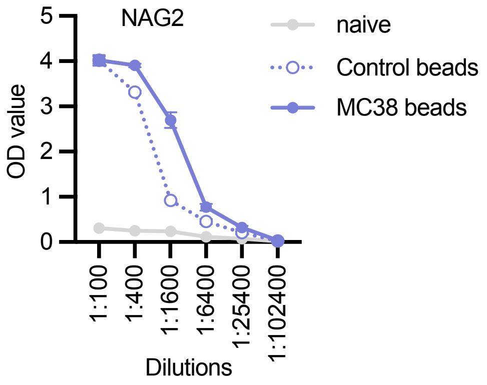


C

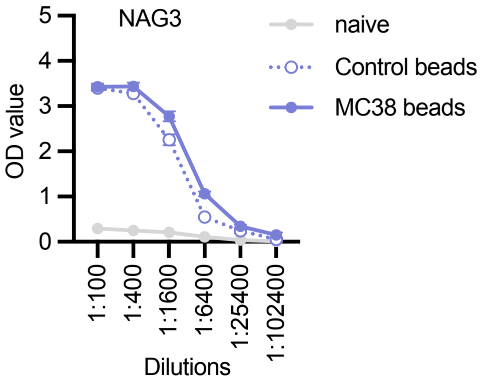


D

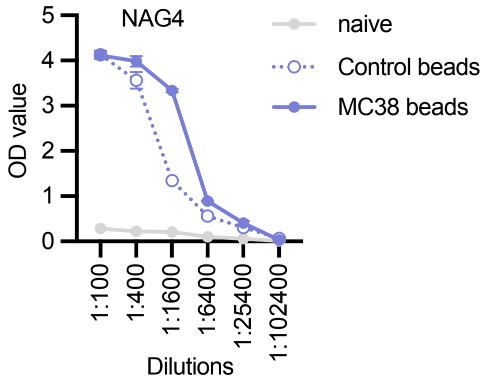


E

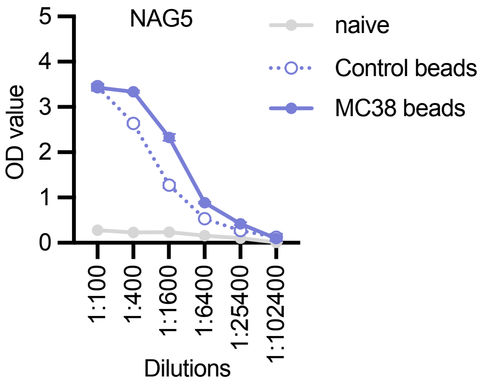


F

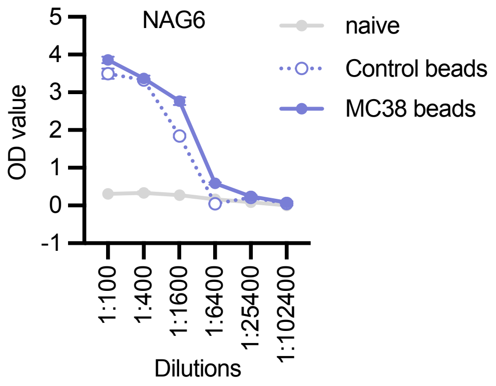


**Supplementary Figure 4.** ELISA data of different sera dilutions
**A-F.** Sera were collected from naïve mice and vaccinated mice (2 weeks after the 3rd vaccination) and were diluted from 1:100 to 1:102400 in ELISA plate coated with NAG1 (**A**), NAG2 (**B**), NAG3 (**C**), NAG4 (**D**), NAG5 (**E**), and NAG6 (**F**) MC38 beads. OD value were measured. Data are shown as mean ±SEM, n=3.

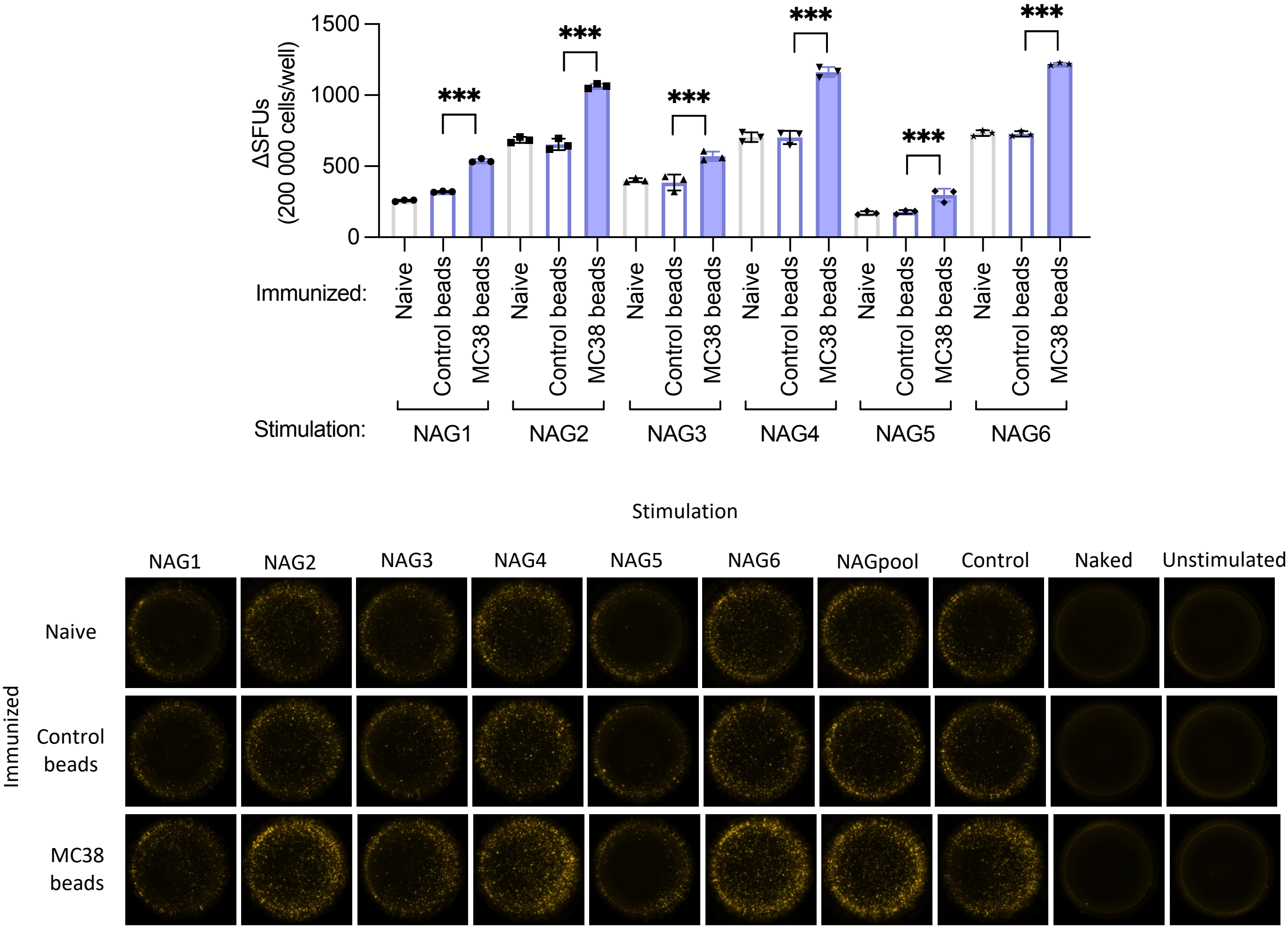


**Supplementary Figure 5.** T cell re-stimulation assay

Splenocytes were isolated from naïve mice and vaccinated mice (2 weeks after the 3rd vaccination) and stimulated with NAG1 to NAG6 MC38 beads. TNF-α FluoroSpot plate were used to detect TNF-α formation. Spot forming units were background adjusted of uncoupled naked (activated/deactivated) bead wells by subtracting the SFUs. Data are shown as individual values, mean ± SEM, n=3. ***, $p<0.001$ by one-way ANOVA and Tukey post-test.

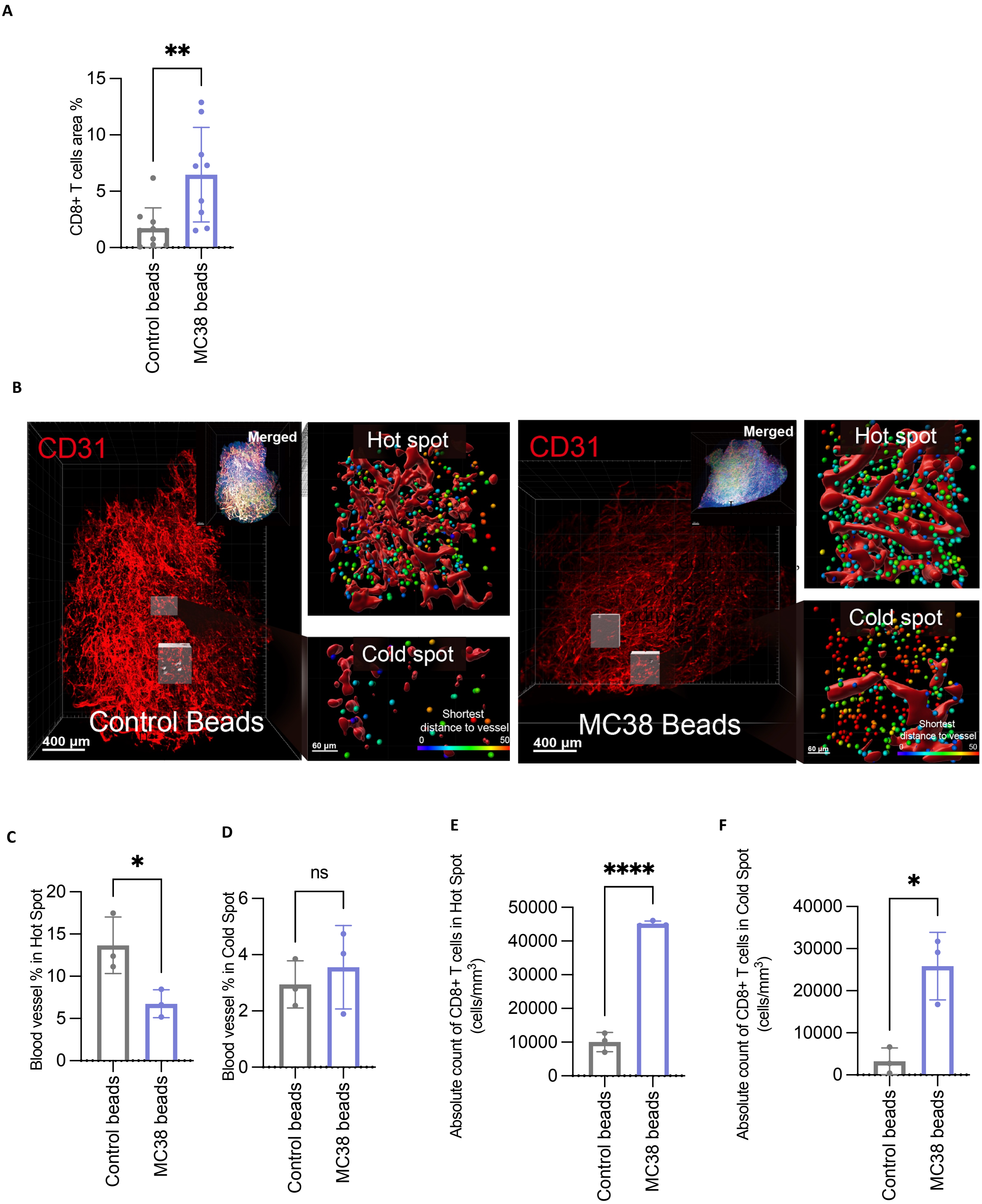


**Supplementary Figure 6.** MC38 beads vaccination can inhibit MC38 tumor growth

**A-F**. Tumor were isolated when the size reached 200-300mm$^3$, and 3D imaging were performed. CD8 T cells were measured (A, E, F). CD31 were stained and blood vessels were measured (B-F). Blood vessels are measured in Hot Spot (C) and Cold Spot (D). CD8 T cells were calculated in Hot Spot (E) and Cold Spot (F). Data are shown as individual values, mean ± SEM, n=9 (A), n=3 (C-F). ns, not significant; *, p<0.05; **, p<0.01; ****, p<0.0001 by unpaired t test (A, C-F).

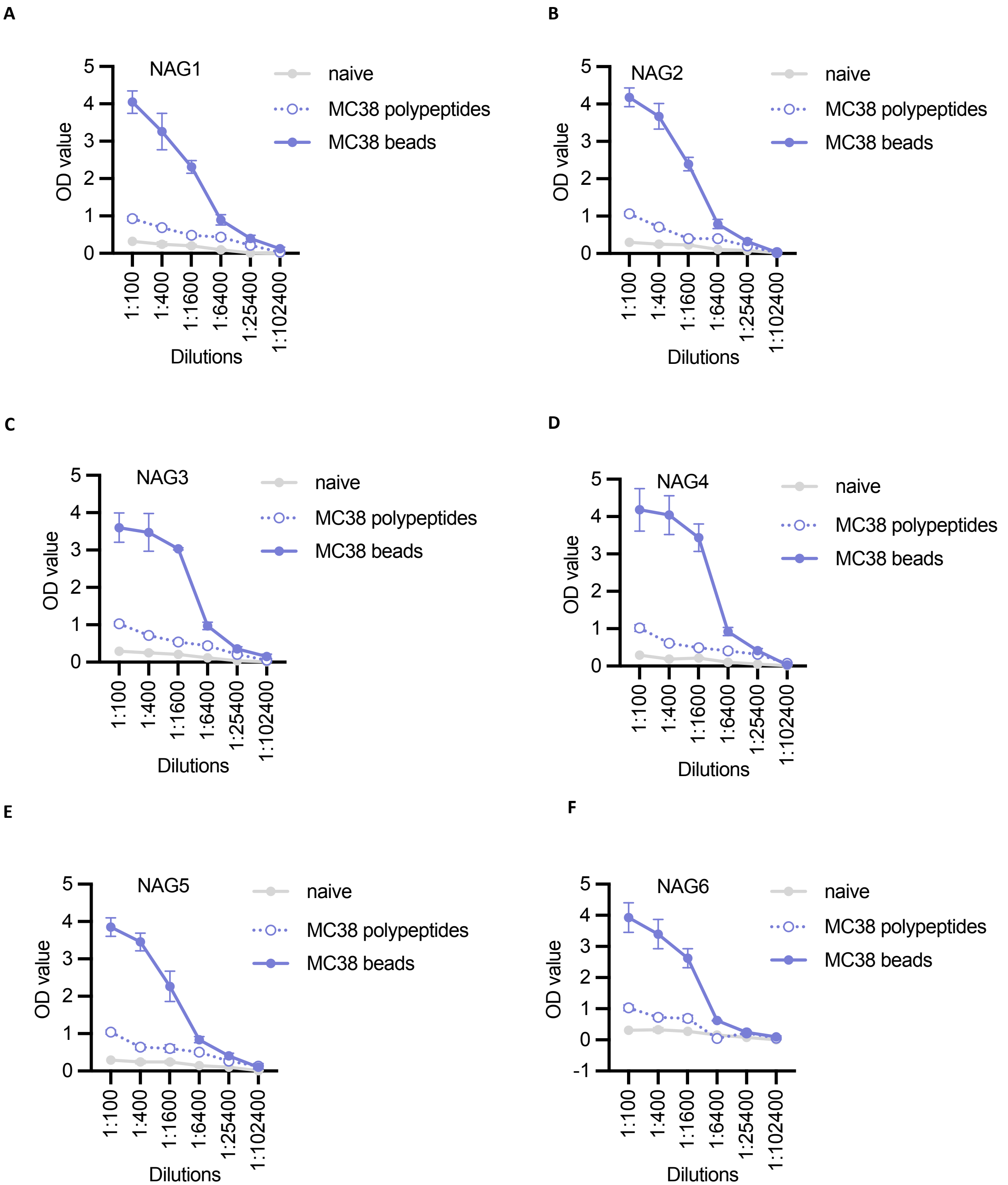


**Supplementary Figure 7.** ELISA data of different sera dilutions

**A-F.** Sera were collected from naïve mice, and 2 weeks after the 3rd polypeptides and beads vaccinated mice and were diluted from 1:100 to 1:102400 in ELISA plate coated with NAG1 (A), NAG2 (B), NAG3 (C), NAG4 (D), NAG5 (E), and NAG6 (F) MC38 beads. OD value were measured. Data are shown as mean ±SEM, n=3.

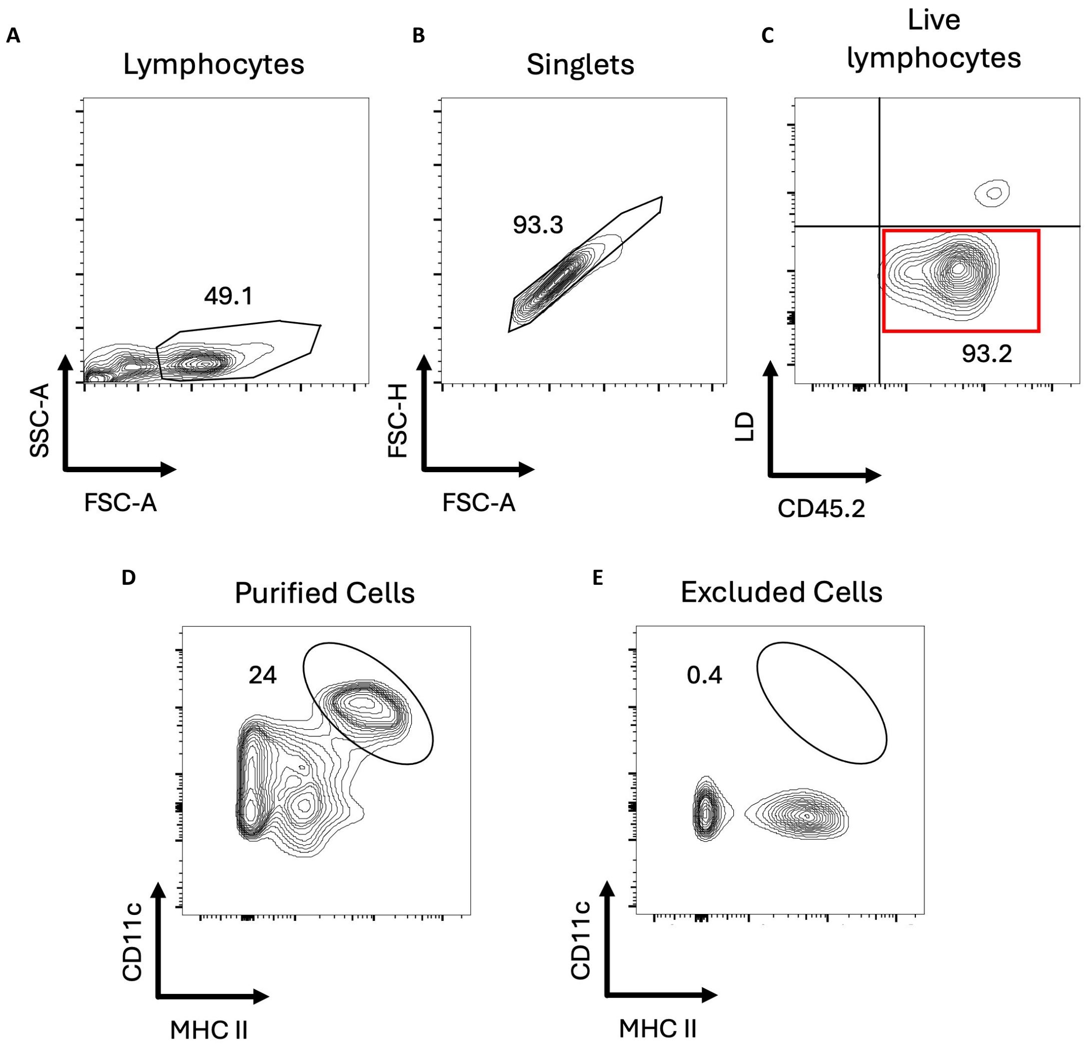


**Supplementary Figure 8.** Dendritic cells isolated from murine spleen and lymph nodes.
**A-C** Gating strategy of cells obtained from dendritic cells isolation. All single cells (A-B) are gated based on CD45+ cells that are alive (**C**). **D-D** Expression of dendritic cell marker on isolated cells. CD11c and MHC class II are used as a marker for dendritic cells. At least 24% of cells that isolated with the pan mouse dendritic isolation kit (Miltenyi) are positive for dendritic cells marker (**D**), compared to 0.4% of cells from the remaining cells excluded during the isolation process (**E**).

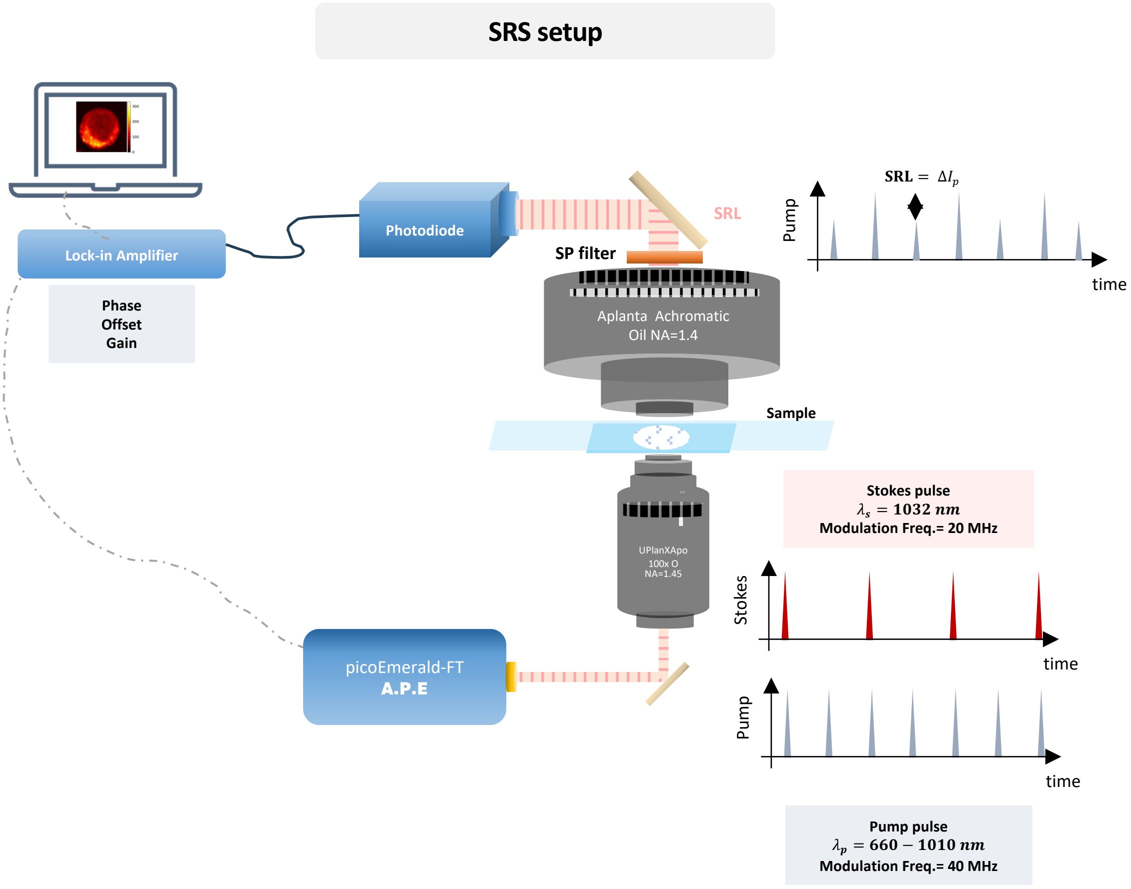


**Supplementary Figure 9.** Schematic of the 3D SRS microscopy setup.

Pulsed pump and Stokes beams from the picoEmerald-FT laser are spatially and temporally overlapped, with the Stokes beam fixed at 1032 nm and the pump beam tuneable from 660–1010 nm. The combined beams are focused into the sample via a 100× oil-immersion objective. After interacting with the sample, the transmitted signal is collected through an oil-immersion condenser and directed to a APE-detection module with a Si-photodiode and lock-in amplifier, synchronized to the laser. This setup enables high-resolution, label-free 3D SRS imaging of cellular structures.

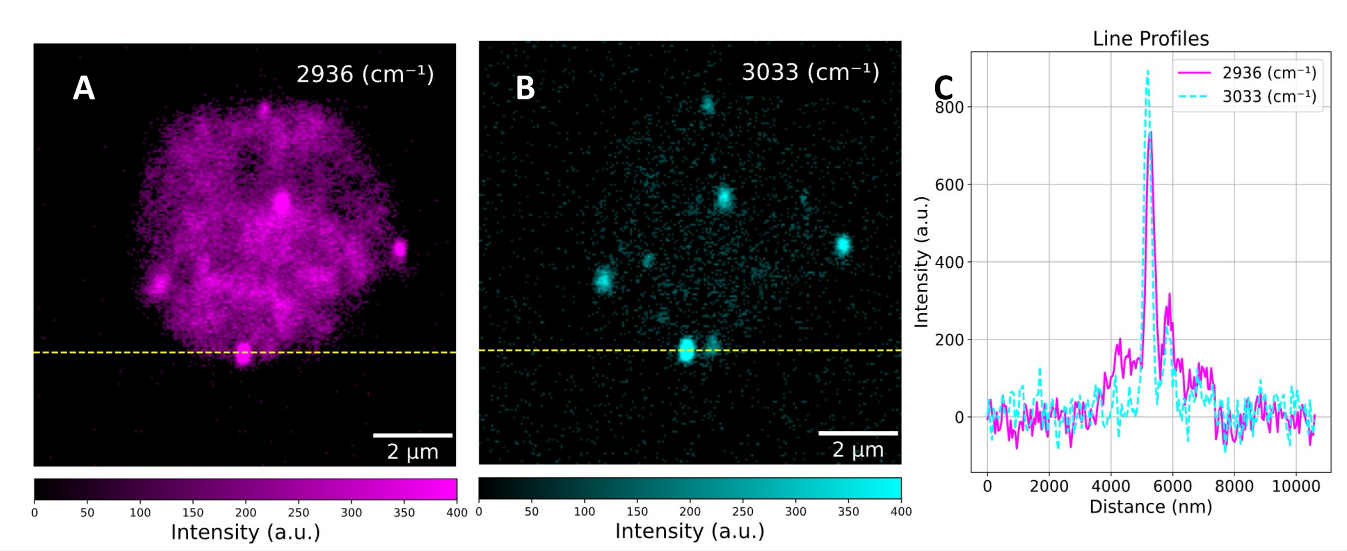


**Supplementary Figure 10.** Visualization and sizing of internalized beads in APCs by 2D SRS.
2D SRS imaging of a representative antigen-presenting cell reveals seven internalized beads. Line profiles of protein (2936 $cm^{-1}$, magenta) and bead (3033 $cm^{-1}$, cyan) signals allow direct estimation of invaded bead's size via the 3033 $cm^{-1}$ FWHM.
