## Supplementary Tables for "A neoantigen-microbead platform for personalized T cell cancer vaccination"

| VarID | Ref | Alt | Score | Gene | Variant (P) | NAG construct # |
|---|---|---|---|---|---|---|
| X:101596244:G:T | G | T | 97 | Taf1 | p.G1824V | 1 |
| 4:138221673:C:T | C | T | 97 | Hp1bp3 | p.L16F | 2 |
| 13:119477808:CA:AG | CA | AG | 91 | 4833420G17Rik | p.T484K | 3 |
| 17:15768733:G:T | G | T | 85 | Chd1 | p.G1583C | 4 |
| 6:31451554:G:T | G | T | 85 | Mkln1 | p.G273C | 5 |
| 18:42333348:G:T | G | T | 81 | Rbm27 | p.G881V | 6 |
| 14:101919281:A:T | A | T | 80 | Lmo7 | p.T1296S | 1 |
| 14:101919443:G:A | G | A | 79 | Lmo7 | p.E1350K | 2 |
| 6:117923784:G:A | G | A | 79 | Hnrnpf | p.G10S | 3 |
| 14:56797820:C:A | C | A | 77 | Zmym5 | p.V264L | 4 |
| 11:101572554:G:C | G | C | 77 | Nbr1 | p.G616R | 5 |
| 9:25130622:C:G | C | G | 76 | Herpud2 | p.V85L | 6 |
| 6:94609026:G:T | G | T | 76 | Lrig1 | p.T727N | 1 |
| 4:107037631:A:G | A | G | 76 | Ssbp3 | p.N249S | 2 |
| 14:56912999:G:T | G | T | 75 | Zmym2 | p.S393I | 3 |
| 4:108642207:T:G | T | G | 73 | Zfyve9 | p.S614R | 4 |
| 6:125101598:G:T | G | T | 72 | Chd4 | p.A228S | 5 |
| 4:129341521:G:C | G | C | 72 | Zbtb8os | p.C56S | 6 |
| 11:88837167:G:T | G | T | 72 | Akap1 | p.T663N | 1 |
| 17:65838926:G:A | G | A | 71 | Ppp4r1 | p.A887T | 2 |
| 11:69589202:G:T | G | T | 71 | Trp53 | p.G239V | 3 |
| 11:59737404:G:T | G | T | 71 | Mprip | p.G226W | 4 |
| 4:101163681:C:A | C | A | 70 | Jak1 | p.G654V | 5 |
| 4:43543211:C:A | C | A | 70 | Tln1 | p.A1318S | 6 |
| 14:20764912:C:A | C | A | 70 | Camk2g | p.W215C | 1 |
| 4:101163722:C:T | C | T | 69 | Jak1 | p.M640I | 2 |
| 11:107052931:C:A | C | A | 69 | Bptf | p.R2593L | 3 |
| 14:25645744:G:T | G | T | 67 | Zmiz1 | p.A282S | 4 |
| 15:82194964:G:T | G | T | 67 | Srebf2 | p.D838Y | 5 |
| 19:41238809:C:A | C | A | 67 | Tm9sf3 | p.Q274H | 6 |
| 17:64292869:C:A | C | A | 66 | Pja2 | p.S540I | 1 |
| 14:70692710:C:A | C | A | 66 | Xpo7 | p.V349L | 2 |
| 3:88700451:G:C | G | C | 66 | Khdc4 | p.G369R | 3 |
| 17:84994265:G:T | G | T | 65 | Ppm1b | p.R191L | 4 |
| 3:19211545:G:T | G | T | 65 | Mtfr1 | p.L81F | 5 |
| 8:84698240:G:A | G | A | 65 | Trmt1 | p.A484T | 6 |

**Supplementary Table 1.** 36 mutations with the highest score identified in MC38 cells.
The mutations were ranked based on the score, which reflects the allele frequency of the mutated gene, its expression level, and whether the gene is related to cancer.

| Beads dilution | 1:100 | 1:1000 | 1:10000 |
|---|---|---|---|
| Day 1 | 4.3±3.6 | 3.4±4.2 | 3.0±3.2 |
| Day 2 | 2.4±2.7 | 3.0±4.1 | 3.4±3.8 |
| Day 3 | 1.9±2.4 | 2.2±2.7 | 3.5±4.9 |

**Supplementary Table 2**. Bead uptake in APCs across concentrations and incubation times. Average number of internalized beads per cell (Mean ± SD) measured from SRS imaging of 450 cells across three incubation durations and multiple bead concentrations.
